# Senescent and reactive astrocytes display distinct expression profiles and divergent functional capacities

**DOI:** 10.64898/2026.06.05.729920

**Authors:** Sabina B. Knox, Luisa M. Abadia, Nicholas J. Guzman, Eishi Noguchi, Qiang Liang, Christian Sell

## Abstract

Astrocytes assume multiple phenotypes in the brain in response to stress, injury, inflammation, and aging. Given the complexity of this critical cell type in the CNS, it is important to gain a greater understanding of the differences between these phenotypes and to potentially identify therapeutic approaches to modifying astrocyte function in the context of disease and aging. We compared senescent and reactive astrocytes using a strictly defined paradigm to induce these phenotypes in human astrocytes. Gene expression profiling reveals overlapping but distinct expression profiles. Reactive astrocytes predominantly express genes involved in inflammatory responses while senescent astrocytes express genes and a secretome that suggests a role in synaptic pruning. Unexpectedly, functional analysis in a simplified neurite outgrowth assay suggests that senescent astrocytes retain the ability to support neurite outgrowth while reactive astrocytes lose this capacity. The data suggests that senescent and reactive astrocytes play distinct functional roles in the physiology of the aging brain. However, the overlapping inflammatory nature of senescent and reactive astrocytes makes it difficult to discriminate between them using existing toolsets designed to identify senescent cells.

## Introduction

Astrocytes are the most abundant glial cell in the central nervous system (CNS) and perform a myriad of functions; forming the blood brain barrier, participating in immune responses, and providing neurotrophic support including neurotransmitter uptake, synaptogenesis, as well as maintaining and pruning synapses (1). Astrocytes support neuronal function through multiple mechanisms including secreted factors, extracellular matrix modulation, and cell surface receptors (2–4). Interestingly, astrocytes provide a complex system that creates restrictive boundaries (5), but also the support for increased dendritic arborizations (4). The mechanisms of neuronal support can also provide cues for synaptic pruning and plasticity (6). In addition to their function in normal brain tissue, astrocytes participate in damage response, repair, and remodeling in pathologic states, following injury and during normative aging (7). The response of astrocytes in these settings is variable and likely influenced by the specific type of damage or insult to the CNS (8). Genetic profiling of astrocytes comparing ischemic stroke and neuroinflammation indicates changes over time in astrocyte response that appears to be specific to the type of insult (9). Thus, astrocytes represent a critical and dynamic cell population within the CNS that is central to neural function. The integration of these mechanisms indicates that astrocytes are vital to the complex formation of neural circuitry in the brain (10).

Following injury, astrocytes undergo reactive astrogliosis, a complex response as part of the neuroinflammatory response (11). The specific response of astrocytes depends on many factors including the type of injury or disease, the stage of pathology, location in the brain, as well as pre-existing heterogeneity (12). In some cases, this complex process has been conceptually reduced to two states: A1 and A2 (12, 13). Although there is debate regarding specifics of the reactive phenotypes, A1 phenotype is considered neurotoxic and commonly follows a neuroinflammatory response to injury while the A2 phenotype is considered neuroprotective and is associated with ischemic injury (14, 15).

Aging in the brain is accompanied by the apparent accumulation of both neural and glial cells that have acquired a senescent phenotype as measured by markers such as the cell cycle inhibitors p21^Cip1^ and p16^INK4a^ (16). Our current understanding of senescence suggests that senescent cells may contribute to functional decline and loss of homeostasis that is characteristic of human aging and that senescent cells in the CNS may contribute to age-related decline in neural function and pathologies such as Alzheimer’s disease, as has been described in other tissues (17). At the fundamental level, cellular senescence is a programed cell response triggered by unresolved damage or chronic stress. Senescent cells display extensive changes in cell metabolism and reorganization of chromatin domains. We have previously characterized replicative senescence in human astrocytes and described a differential sensitivity to oxidative stress relative to human fibroblasts (18). In addition, we have identified increased numbers of astrocytes harboring markers of senescence in human brain tissue from patients with Alzheimer’s disease and aged brains (19), and these cells appear to contribute to cognitive decline (20). Other studies have shown that astrocytes undergo senescence in response to irradiation (21) and clearance of senescent glial cells reduces tau-dependent pathology (22).

The relationship of senescence in astrocytes relative to their functional activation during response to damage, injury, or infection is of critical interest as it will be important to understand how potential therapeutic targeting of senescent astrocytes will intersect with their function in the CNS. Cytokines that trigger reactive astrocytes have been defined in the context of LPS challenge in mice (23, 24) and targeting these cells appears to be beneficial in the context of Amyotrophic lateral sclerosis (25). Comparison of human astrocytes forced into a reactive state using these cytokines display some overlap with H_2_O_2_-induced senescence (26), however the precise relationship between activated astrocytes and senescent astrocytes remains unclear.

In this work, we examine the potential for endogenous replication and transcriptional stress to induce senescence in human astrocytes. We develop a model system to study senescence in human astrocytes in a controlled manner and examine the relationship of cells that enter senescence to reactive astrocytes.

## Results

We considered the potential sources of endogenous stress and damage that cells of the CNS may face including oxidative stress, replication stress, and transcriptional stress among others. Of these, the most quantifiable are replication and transcriptional stress as immunostaining can reveal foci of unrepaired DNA damage. Our objective was to create a model that can be used to evaluate the response to unresolved damage across cells of the CNS. Therefore, focused on transcriptional stress as both post mitotic neurons and glial cells may be subject to endogenous transcriptional stress. Because topoisomerase II is essential for the resolution of DNA breaks caused by transcriptional stress, we evaluated the ability of topoisomerase II inhibition to induce senescence in human primary human astrocyte cultures. To reduce the impact of replicative stress, we used a 30-minute window of exposure to the topoisomerase II inhibitor etoposide (Fig 1A) and evaluated CDKN1A and LMNB1 mRNA levels at timepoints following damage and titrated the concentration of ETO to produce a treatment that would induce senescence at the lowest possible level of persistent damage. Interestingly we found that human astrocytes are much more resistant to etoposide than human neuronal cells. Two 30-minute exposures using 100 μM ETO were required to produce a low level of persistent DNA damage as assessed by γH2AX and p53BP1 foci. This exposure produced 1-5 foci per cell at 7 days post damage. This concentration and time of exposure was evaluated in more detail because human cells have high fidelity genomic surveillance and 1-2 sites of unresolved damage are sufficient to induce senescence (27). A significant increase in CDKN1A and a decrease in LMNB1 mRNA levels (Fig 1B-D) further confirmed that this level of DNA damage would trigger the senescence response in these cells. As a further measure of senescence, we examined SA-βGal staining and there were significantly more SA-βGal positive cells in the senescent group than the control (Fig S1). These results verify that this low-level damage regimen can reliably induce senescence in primary human astrocytes.

**Figure 1:**
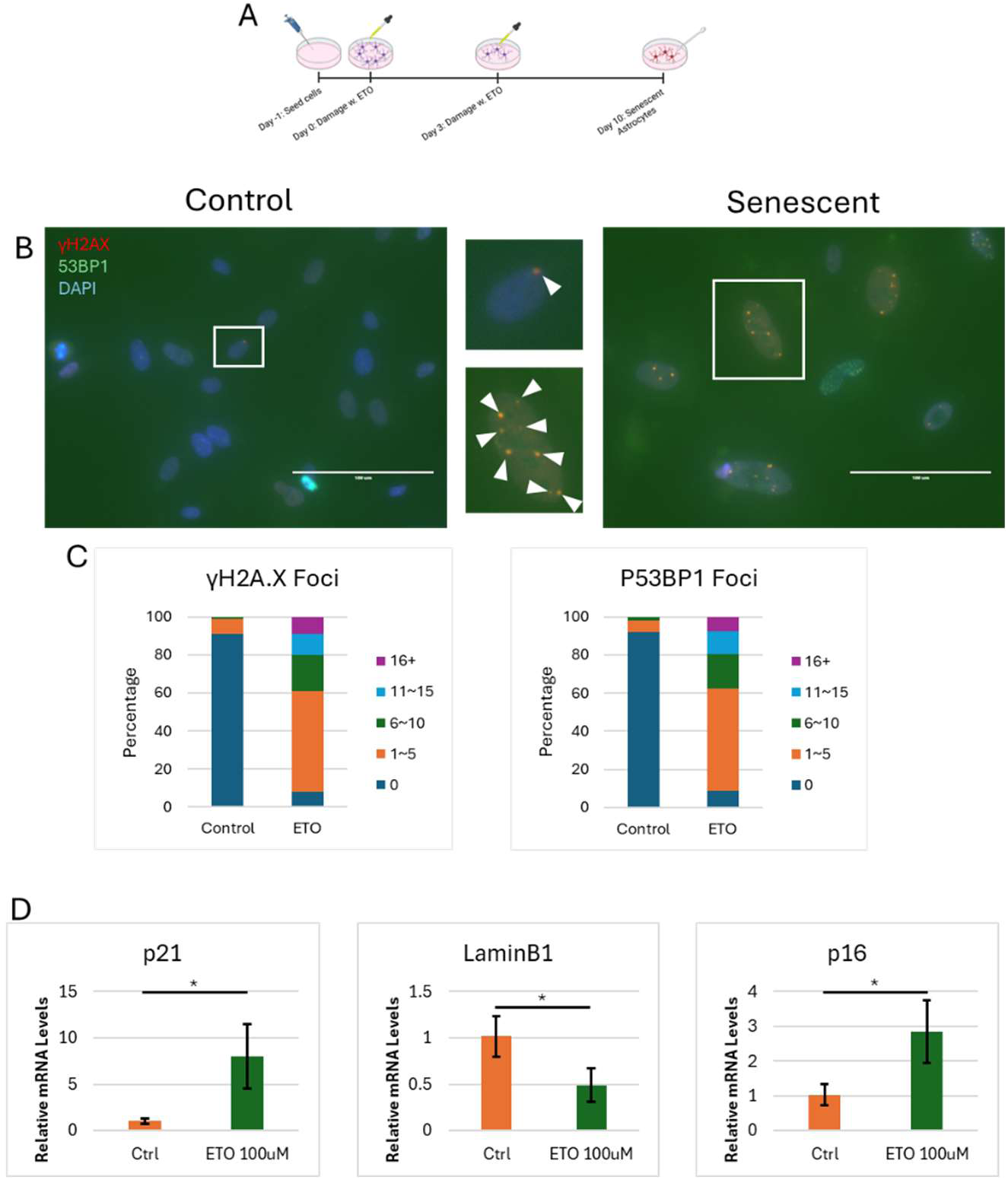
Human astrocytes display senescent features following exposure to etoposide. Panel A contains a schematic of the senescence induction protocol which involved 2 30 minute exposures to 100μM etoposide separated by 3 days and followed by a 10-day recovery period to allow full expression of the senescence program. Panel B contains representative images of immunofluorescence imaging of control and senescent astrocytes displaying γH2AX and 53BP1 localization. Panel C contains quantification of γH2AX and p53BP1 immunofluorescent images using ImageJ software, comparing percentage of cells in each group of number of puncta per cell. astrocytes 10 days following the second damage. Panel D contains graphs of the relative mRNA expression from RT-qPCR of p21 (CDKN1A), LaminB1 (LMNB1), and p16 (CDKN2A). Data are represented by mean ±SD. Statistical analysis was performed using a two-tailed unpaired student’s t-test, with significance levels marked as *p < 0.05, **p < 0.01, ***p < 0.001, and ****p < 0.0001.

To further characterize the senescent phenotype in human astrocytes we performed bulk RNA-seq. We found that the senescent cells had a distinct transcriptomic profile from the control (Fig 2A, S2A) with approximately 400 genes upregulated and downregulated. As expected, most genes that were downregulated are involved in cell-cycle progression; conversely, most genes that were upregulated are involved in inflammatory pathways (Fig S2B). Individually, the most upregulated genes were SOD2, CDKN1A, C3, IL6, and IL8. SOD2 has been strongly associated with age-related decline in the nervous system and specific targeting of SOD2 in astrocytes either reducing or increasing SOD2 has profound impact on cognitive function in mice (28, 29) (30) while deletion produces a progeroid syndrome (31). CDKN1A confirms that our protocol induces key mediators of the senescence program while IL6, and IL8 are classic features of senescence associated with cell cycle arrest and SASP, respectively (34). C3 is known to function in the development of the nervous system, mediating synaptic plasticity and pruning (32, 33), proving an interesting feature of aging and possibly a factor in age-related cognitive decline. Gene ontology analysis revealed a downregulation of cell cycle associated pathways (G2M, E2F, and Myc targets) concomitant with an upregulation of the p53 pathway, TNF α, and IL6 STAT3 pathways (Fig S2B).

**Figure 2:**
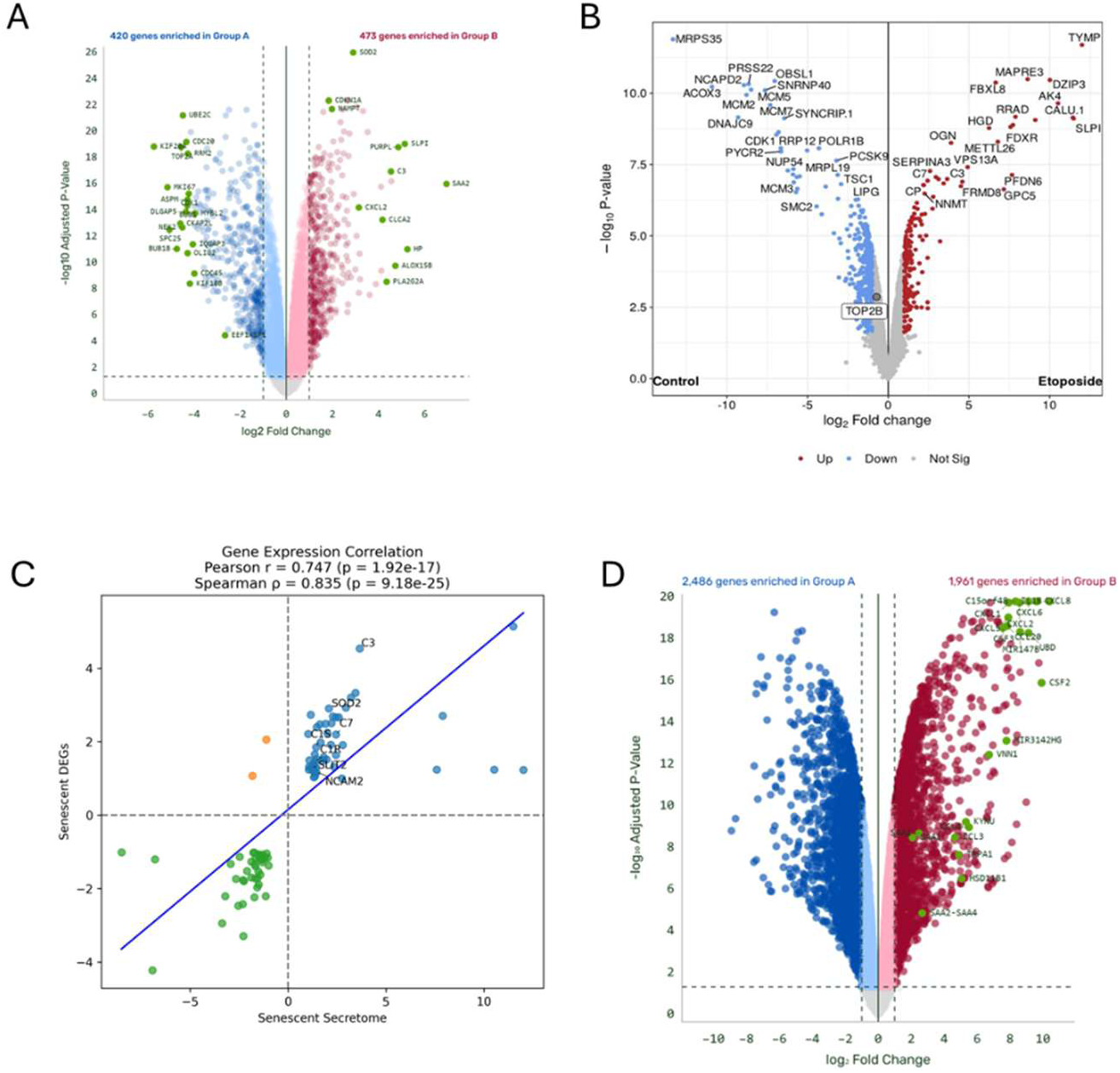
Senescent astrocytes display a distinct transcriptional and secretory profile. Volcano plots of RNA-seq transcriptional analysis (Panel A) and secreted proteins (Panel B) for senescent (ETO treated). Panel B contains a volcano plot of secretory proteomics analysis from senescent astrocytes. Panel C contains a correlation analysis of overlapping genes between RNA-seq data from A and secretory results in B. Blue: Up/Up, Orange: Down/Up, Green: Down/Down, and Red: Up/Down, respectively. Statistical correlation was calculated utilizing both Pearson and Spearman analyses. Panel D contains a volcano plot comparing cytokine-treated, reactive astrocytes (Group B-red) and control cultures (Group A-blue).

We also explored the secretory phenotype of the senescent astrocytes to compare transcriptional and proteomic changes. As expected, the secretory phenotype of the senescent cells proved to be distinct from the control (Fig 2B). Additionally, the proteome of the secreted factors significantly aligned with the transcriptome with an overlap of ∼90 genes/proteins (Fig 2C). However, against expectations, there was no significant difference in IL6 and IL8 levels (Fig 2B), two prominent SASP factors (34). The lack of IL6 may be due to the limited sensitivity of the proteomic analysis however; the secreted inflammatory factors highly correlated with transcriptional changes and components of the complement system involved in synaptic pruning such as C3, C7, C1s, and C1r (33) correlated well (Fig 2C). Furthermore, there was an upregulation in factors involved in neuronal migration such as SLIT1(35) and NCAM2(36) (Fig 3B), indicating there may be more neuronal support provided than previously theorized for senescent astrocytes. Together, these results show that senescent astrocytes display a distinct transcriptional and translational shift from control astrocytes but may not align with the conventional SASP function in senescent cells from other tissue types.

**Figure 3:**
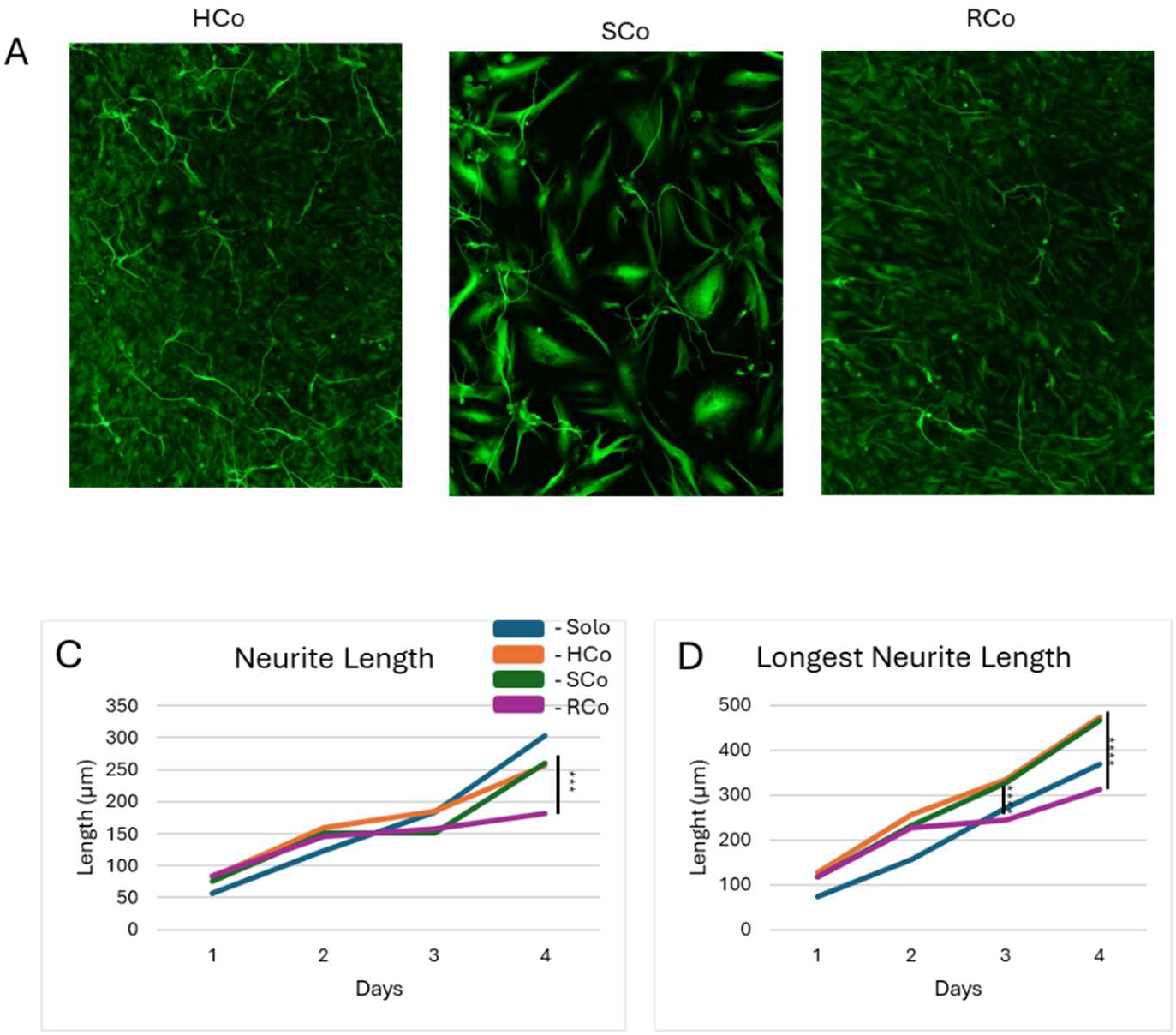
Senescent astrocytes support iNeuron outgrowth while reactive astrocytes are impaired. A) Representative images of solo, HCo, senescent SCo, and reactive RCo astrocyte/neuron co-cultures at 20x. B) Median length of all analyzed neurites over 4 days. C) Median length of longest neurite per cell over 4 days. Statistical analysis for neutite measurements were performed using a two-tailed unpaired student’s t-test and Mann-Whitney test with significance levels marked as *p < 0.05, **p < 0.01, ***p < 0.001, and ****p < 0.0001.

To gain insight into the distinction between senescent astrocytes and reactive astrocytes, we treated human astrocytes with Il-1α, TNFα, and C1q, cytokines produced endogenously by microglia That have been shown to produce the A1 neurotoxic-like reactive phenotype (24). 24 hours post treatment the transcriptome of these cells was evaluated in comparison to the untreated control cultures. This comparison revealed a larger divergence in gene expression than the senescent comparison. A total of 2486 genes were downregulated and 1961 were significantly upregulated when comparing control and reactive at 24 hours (Fig 2D). Gene ontology analysis revealed upregulation of pathways associated with proliferation such as E2F, G2M, and Myc targets in contrast to the senescent astrocytes. However, inflammatory pathways such as TNF α, Interferon γ, and IL6 STAT3 were upregulated as in senescent astrocytes (Fig S3A). We used GSEA analysis to compare the senescent astrocyte signature to the reactive astrocyte signature in key pathways. GSEA analysis for astrocyte activation revealed the expected significant increase relative to control (Fig S3B). GSEA specific to the key inflammatory response from NFκB was significantly increased in both senescent and reactive gene sets (Fig S3C). Because both the reactive and senescent GO analyses flagged the complement pathway, we looked at the complement cascade activation and found that both phenotypes show significant upregulation of the complement cascade (Fig S3C).

The A1 neurotoxic reactive astrocyte secretome has been reported to include an upregulation of cytokines that aligned with classic SASP markers(14); therefore, we examined the difference in the SASP transcriptomes. Both the senescent and reactive astrocytes significantly upregulate this pathway (Fig S3C). To further investigate the differences between the two phenotypes, we examined the fold change for specific genes involved in all three upregulated pathways. The NF-κB and SASP pathways overlap in genes such as IL6, 8, and 11 as well as CXCL2 and 6. As expected, both phenotypes display a significant increase in fold change, with the exception of IL11 in senescent cells (Fig S3E). It is notable that the reactive phenotype displays a much larger difference in fold change to the control compared to the senescent phenotype in the SASP markers (Fig S3E). Interestingly, the complement factors C3, C1R, and C1S followed a slightly different pattern. The senescent cultures displayed a significant increase in all three factors, whereas the reactive cultures significantly increased for C3 and C1S but not C1R (Fig S3E). Overall, these results show that both the reactive and senescent transcriptomes both contain elevated levels of mRNAs coding for inflammatory cytokines but reactive astrocytes have a stronger inflammatory profile while senescent astrocytes produce higher levels of several complement components associated with synaptic pruning.

Because senescent astrocytes have been likened to reactive astrocytes that do not divide, we sought to analyze their transcriptomes in cell division pathways. We analyzed the transcriptional changes in the E2F and G2/M pathways which are involved in cell cycle progression and division. Consistent with enhanced proliferation, reactive astrocytes displayed a significant upregulation of these pathways relative to control cultures while senescent astrocytes showed a significant downregulation (Fig S3F) further emphasizing the distinct difference in cellular division. To provide a global comparison of the two transcriptomes we directly compared the most significantly upregulated and downregulated genes. Using stringent cutoff markers, we narrowed the list of DEGs to genes that are both significant and have a │log_2_│≥ 1. Next, we found about 500 overlapping DEGs between the two phenotypes and compared their expression patterns. Based on our previous results, we expected there to be a significant correlation between the two phenotypes. However, in an overall comparison for the expression profiles the two transcriptomes proved to be uncorrelated (Fig S3G). Thus, there are distinct and significant differences between reactive and senescent astrocytes that remain to be defined.

In our secretome analysis, we observed an upregulation of factors that impact specific aspects of astrocyte function such as the guidance factors SLIT2 and NCAM2(35–37). A neuron outgrowth assay allows functional analysis of senescent astrocytes that may reflect differences related to the transcriptome. We utilized hiPSC-derived neurons (iNeurons) cocultured with control (HCo), senescent (SCo), or reactive astrocytes (RCo) to evaluate the impact of different astrocyte phenotypes on neurite outgrowth (Fig 3A). Interestingly, there was a significant decrease in both neurite length and longest neurite length in cultures containing reactive astrocytes (Fig 3 B, C) but there was no significant difference between the healthy coculture, the senescent coculture, or the reactive coculture in terms of the number of neurites per cell (Fig S4A). However, all three conditions were significantly higher than the iNeuron-only (Solo) culture (Fig S4A). To integrate these functional differences into the transcriptional data, we compared gene sets related to neurite growth pathways. We found that reactive astrocytes significantly downregulate axon guidance and NCAM signaling pathways whereas senescent astrocytes showed no significant shift in these pathways (Fig S4B).

We examined the changes in specific genes within the datasets which are involved in the overall process. Consistent with the GSEAs, there was a significant reduction in genes such as NCAM1 and NCAM2 as well as SARC and SPARCL1 in the reactive astrocytes (Fig S4E). The senescent gene profile showed increased expression of these genes except for CSPG4 (Fig S4E). These results suggest that neurite outgrowth is a distinction between the two phenotypes and further emphasize the difference seen in their transcriptomes.

It has been suggested that reactive astrocytes may progress to senescence over time(38) and our in vitro model allows us to test this directly by incubating cytokine treated astrocytes for extended periods. We maintained reactive astrocytes for 14 days and examined the transcriptome at 7 and 14 days (Fig S5A, B). Geen ontology analysis indicates that the reactive cultures were relatively enriched in proliferation associated mRNAs at day 1 but were more enriched in inflammatory mRNAs relative to controls at days 7 and 14 (S5B). There were significant changes in the gene expression patterns and a progressive increase in inflammatory cytokines such as CXCL4, 6, 8 and 20. The mRNA levels for serum amyloid proteins A1, A2, and A4 increased relative to control over the course of the 7 and 14 day points while IL1β mRNA was relatively elevated at day 1 but much less so at days 7 and 14(Fig S5C). Interestingly there was no increase in CDKN1A (p21) indicating that there is no transition in cell cycle arrest, suggesting that the reactive astrocytes maintain their inflammatory profile without a transition towards a senescent state.

Given the strong inflammatory profile of both the senescent and reactive astrocytes, and the fact that the gene profiling data sets for senescence (SenMayo, Fridman) contain multiple factors common to both phenotypes such as C3, IL8, and TNF responsive genes. We compared the gene expression profiles to these gene sets using GSEA. The senescent astrocyte transcriptome significantly aligned with established senescent sets (both Friedman and SenMayo gene sets) as well as the p53 signaling pathway, verifying their senescent identity (Fig 4A). These results highlight the altered transcriptional profile and shift in molecular pathways in senescent astrocytes. Interestingly, we find that the cytokine treated, reactive astrocytes scored significantly in both the Fridman and SenMayo senescence gene profiles (Fig 4A), potentially due to the shared inflammatory profile.

**Figure 4:**
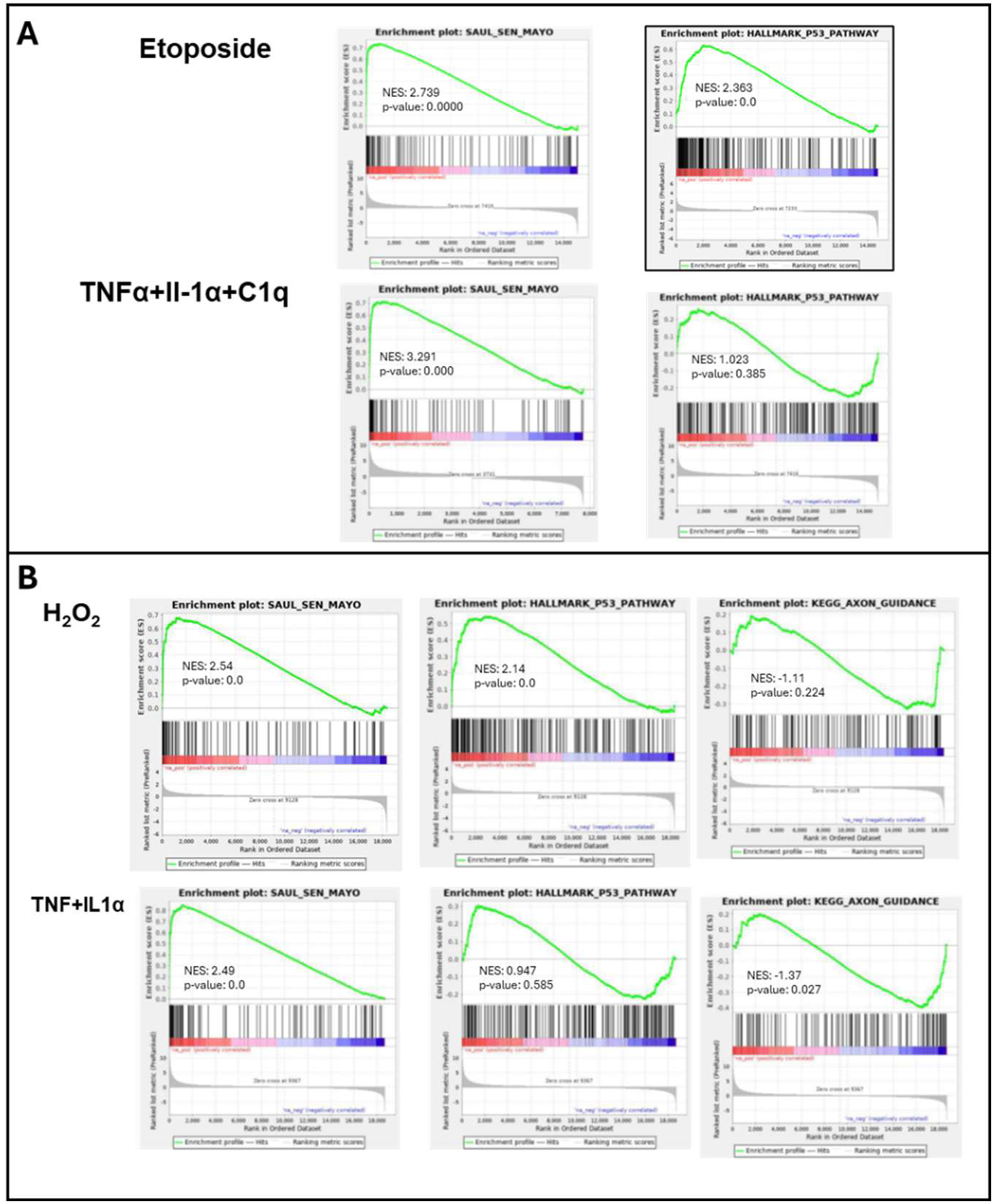
Expression profile differences between senescent and reactive astrocytes are similar in published data sets. Differential gene analysis performed on data from Simmnacher et al 2020(26) was used for GSEA analysis using a senescence gene set (SenMayo), a p53 response gene set, and an axonal guidance gene set. Senescent cultures (H_2_O_2_) were significantly different from control for SenMayo and the Hallmark p53 pathway but not for Axonal guidance while the reactive culture expression profile (TNF + IL1α) were significant for SenMayo and axonal guidance but not for the Hallmark p53 pathway.

In addition, we examined a published data set from the Winner laboratory which utilized the same commercial source of astrocytes and followed an H_2_O_2_ stress protocol developed in our group to examine the difference between human fibroblasts and astrocytes(18, 26). In this case, human astrocytes were induced to senesce using H_2_O_2_ and activation induced by the combination of TNFα and IL1α. Interestingly, the gene set analysis showed the same pattern observed in our data set (Fig 4B). All data sets score positive for senescence using the senescence gene sets but only the senescent cultures (ETO and H_2_O_2_ treated) scored positive for p53 activation.

In terms of axonal guidance, the Winner reactive astrocyte data (TNF + IL1α) set also showed a significant decrease in genes associated with axon guidance while the senescent, H_2_O_2_, data set showed no significant difference, mirroring our results. This confirms the gene pattern using an independent data set produced through an alternative approach to the induction of senescence and a slightly different combination of cytokines to produce reactive astrocytes.

## Discussion

We provide new information regarding the senescence program in human astrocytes using a well-controlled approach to produce low levels of persistent DNA damage. The results indicate that the transcriptional and functional properties of senescent and reactive astrocytes are distinct but have multiple common elements. Critically, the existing GSEA profiling gene sets scored both senescent and reactive astrocytes as highly significantly enriched for senescence-associated genes. The ability to identify senescent cells in multiple tissues has been the subject of focused effort in the field and while there has been progress in multiple tissue types (39), the brain lags in terms of both a general understanding of the senescence program and the impact of senescent cells in aging and disease progression. The strong signal from both senescent and cytokine treated astrocytes uncovered in our data underscores the need for a more detailed understanding of senescence and suggests that caution is required when interpreting data from transcriptome profiles for senescence.

Although it may be expected that senescent astrocytes would lack functionality and in some experimental paradigms provide a detrimental environment (40), it appears that the senescent astrocytes were competent in terms of neurite outgrowth in the specific experimental paradigm used in this study. This is a specific setting utilizing iPS-derived immature neurons which are likely to be quite robust in terms of their response to paracrine signaling and outgrowth. It may be that in a setting of neuronal damage the functionality of the senescent astrocytes may differ. However, the data does indicate that the senescent astrocytes provide a superior environment for neurite outgrowth compared to the cytokine-treated, reactive astrocytes. This supports the concept that reactive and senescent astrocytes play distinct roles in the brain during injury and aging. This conclusion is supported by the gene profile analysis indicating that reactive astrocytes have reduced levels of mRNAs associated with axonal guidance and is strengthened by the fact that the data are reproducible from an independent group. Just as the expression profiles of the senescent and reactive astrocytes overlap, one might expect that the functional roles of senescent and reactive astrocytes may be overlapping and complementary to support neuronal function. A detailed examination of the axonal guidance genes that were decreased in the reactive astrocytes reveals multiple members of the semaphoring family as drivers of the decreased axon guidance GSEA in the reactive astrocytes including semphorins 3B, 3E 4B, 4F, 5A, 5G, 6B, and 6D, which are known to act as sell surface guidance factors for neurite outgrowth (41, 42) and apical guidance in neural progenitor cells(43). It may be that the reactive astrocytes produce a negative environment for outgrowth that is not part of the function for the senescent astrocytes.

Reactive astrocytes express a strong inflammatory profile that includes IL6, IL8, IL11, as well as CXCL2 and CCL6. In contrast, senescent astrocytes express these cytokines at much lower levels than the reactive astrocytes while expressing much higher levels of complement factors including C3, C1S, and C1R. In the CNS, the complement system has been shown to mediate synaptic plasticity and pruning (32, 33), but the function of complement components in terms of disease progression and aging is complex (44). Global reduction of C3 has been reported to improve cognitive function in aged mice (45, 46) and inhibition of the C3a receptor has been reported to reduce tau phosphorylation and ameliorate cognitive defects in a mouse model of tauopathy (47). However, results from a C3aR null mouse model suggest a receptor independent pathway (48) indicating a complex relationship. In terms of the senescent phenotype, the expression of several complement proteins suggest that senescent astrocytes may participate in synaptic remodeling.

In summary, we have undertaken a comparison of senescent and reactive astrocytes using a quantifiable approach to the induction of persistent DNA damage and have demonstrated that astrocytes trigger what appears to be a classic senescence program that is characterized by a cell type specific gene expression profile that includes increased expression of genes associated with synaptic pruning and inflammatory responses. Phenotypic and functional differences exist between senescent and reactive astrocytes that should be further examined to understand the impact of these astrocyte specific cell states and to uncover therapeutic approaches to modify the function of both senescent and reactive astrocytes. Modifying the senescent and reactive phenotypes are highly likely to improve outcomes in age-related cognitive decline and the ability to distinguish between these two important cell states is critical to our understanding of age-related changes in the brain.

## Materials and Methods

### Cell Culture

Primary human astrocytes were obtained from ScienCell (Cat. #1800, Lot#41751, 21-week gestation fetus/male). Astrocytes were grown in ScienCell’s Astrocyte Medium (Cat. #1801) with 2% fetal bovine serum (FBS, Cat. No. 0010) and 1% astrocyte growth supplement (AGS, Cat. No. 1852). They were plated and cultured on 50µg/ml PDL-coated plates. hiPSC-derived neurons were developed using a doxycycline-inducible lentiviral construct containing neurogenin-2(NGN2) and Achaete-scute homolog 1(Ascl1) (Addgene #127289) transduced into Kolf2.1J (JIPSC001000) from The Jackson Laboratory plated on 5 µg/cm^2^ synthemax(Corning, Ref.# 3535)-coated plates. Transduced cells were then selected with puromycin and individual clones tested for uniformity in differentiation in response to doxycyline. Clone 8 was chosen based on this process and cultured to ∼80% confluency in StemFlex (Gibco, Ref# A33493-01) media. On Day 0, the media was then replaced with StemFlex media with 10µg/ml doxycycline (Enzo, Cat.#ALX-380-273). On Day 2, the media was then changed to induction media containing 50% Neurobasal media (Gibco, Ref.# 21103-049), 50% StemFlex media, 1X N-2(Gibco, Cat# 17502048) supplement, 1X B27 (Gibco, Cat# 17504044) supplement, 0.5µg/ml Laminin (Reprocell, Cat.# NP892-021), 10ng/ml BDNF, and 10µg/ml doxycycline. On Day 3, the cells were then dissociated with Accutase and plated onto 50µg/ml PDL and 5µg/ml Laminin-coated plates with week 1 maturation media containing Neurobasal media, 10ng/ml NT-3 (Raybiotech, Cat.# 04G0713W), 1X B27 supplement, 0.5µg/ml Laminin, 10ng/ml BDNF, 10µg/ml doxycycline, and 10µM EdU. A half media change was performed on Day 6. On Day 10, a half media change was performed using BrainPhys™ Neuronal Medium (STEMCELL, Cat. #05790) supplemented with 2% NeuroCult™ SM1 Neuronal Supplement (Cat. #05711) and 1% N2 Supplement-A (Cat. #07152). A half media change was performed every 3-4 days for the duration of culture.

### Senescence induction

Etoposide stock was reconstituted to 10mM in DMSO. Astrocytes were cultured to 70-80% confluency. On Day 0 the media was aspirated and changed to Astrocyte Medium with 100µM Etoposide. Cells incubated for 30 minutes in 37℃ before aspirating the media and replacing with Astrocyte Medium. On Day 3, the process was repeated. On Day 10, the cells were considered senescent.

### Cytokine treatment

A1 reactivity was induced following the Liddelow et al. 2017[39] protocol. Astrocytes were grown to 70-80% confluency before being treated with 3ng/ml Il-1α (Peprotech, Cat.# 200-01A), 30ng/ml TNFα (Acro, Cat.# TNA-H4211), and 400ng/ml C1q (Millipore, Cat.# 204876) for 24 hours. Media was then aspirated and replaced with Astrocyte Medium.

### Quantitative Real-Time PCR

RNA was isolated using TRIzol reagent (Thermo Fisher Scientific) and quantified using a NanoDrop 200 spectrophotometer. Equal amounts of total RNA from each sample were analyzed using quantitative real-time PCR using TaqMan Assays (Thermo Fisher Scientific) (GAPDH: Hs02786624_g1; CDKN1A: Hs00355782_m1; LMNB1: Hs01059205_m1; CDKN2A: Hs00923894_m1) run on an ABI™ 7500 Real-Time system. All the samples were done in triplicates. The fold change was calculated using raw data using the ΔΔCT method. GAPDH was used as internal control.

### SA-βGalactosidase Staining

Cells were washed twice with PBS and then fixed with 3% formaldehyde for 3 min at room temperature. Cells were then washed twice with PBS and stained with X-gal (Tenova X1220) (1 mg/mL of X-gal in 40 mM citric acid/Na2HPO4 [pH 6], 5 mM potassium ferrocyanide, 5 mM potassium ferricyanide, 150 mM NaCl, 2 mM MgCl2) for 18 h at 37°C. Cells were washed with PBS twice and imaged using EVOS FL Auto microscope (ThermoFisher). The percentage of positively stained cells was determined by counting seven random fields. Images of representative fields were captured under 40X magnification.

### RNA sequencing

RNA was isolated using TRIzol reagent (Thermo Fisher Scientific) and quantified using a NanoDrop 1000 spectrophotometer. RNA was cleaned using RNA Clean & Concentrator™-25 (Zymo Research, Cat. # R1017). RNA was then aliquoted into dilutions of 30ng/µl and sent to Plasmidsaurus for sequencing. Biological triplicates in technical replicates were used for Control vs Senescent samples and biological singlicate in technical triplicates were used for Control vs A1 Reactive.

Differential gene expression analysis of expression data from Simmnacher et al. we anayzed using GEO2R.

### GSEA analysis

RNA-seq differentially expressed genes were ordered in a rank file format. The ranked format was then input into GenePattern.org [40] through their GSEAPreranked module and compared with human gene sets from GSEA[41, 42].

### Proteomics

Cells were grown to 70-80% confluency in a 10cm dish coated with PDL in Astrocyte Medium with 2% FBS and 1% AGS. Plates were washed once with 1X PBS then five times with serum-free phenol red-free Astrocyte Medium with 1% AGS and the solution was aspirated after each wash. Cells were then incubated for 48 hours in serum-free phenol red-free Astrocyte Medium with 1% AGS. Conditioned media was collected and centrifuged at 500 x g for 5 minutes to remove dead cells and cell debris. The media was then centrifuged through a 0.22µm filter. The media was then supplemented with protease inhibitors 150 µM PMSF (BACHEM, Art. #4000472.0025), 1µg/ml Pepstatin A (RPI, CAS# 26305-03-3), and 1µg/ml Leupeptin (RPI, CAS# 103476-89-7). It was then divided into aliquots of 4-ml before freezing at -80℃ before being sent to the Wistar Institute for analysis. Samples were in biological triplicates and results were normalized to cell count.

### Astrocyte-Neuron Coculture

Astrocytes were plated on PDL and Laminin-coated plates. Senescent and A1 reactive astrocytes were prepared according to their previous protocols. On Day 9 of the senescence induction protocol, the A1 protocol began. On Day 10 of the senescence induction protocol, Day 3 dissociated iNeurons were seeded on the astrocytes in week 1 maturation media. After 4 days in coculture, the media was changed. After 3 more days the media was changed to BrainPhys™ Neuronal Medium.

### Immunofluorescence

Astrocytes and astrocyte-iNeuron cocultures were washed once with 1X PBS and placed on shaker (The Belly Dancer®, Life Science Incorporated) for 5 minutes. Cells were fixed with 4% paraformaldehyde (Biotium, Lot: 21P0610) and then permeabilized in 0.1% Triton X-100 in PBS for 10 minutes. Cells were blocked with 1% Bovine Serum Albumin (BSA-ASH, RMBI incorporated), 2.25% Glycine, and 0.1% Tween-20 in PBS at room temperature for 1 hour. Cells were then incubated with primary antibodies in the blocking buffer overnight at 4°C. Primary antibodies were removed 24 hours later followed by three washes with 1X PBS 5 minutes each. Samples were then incubated with secondary antibodies in blocking buffer for 1 hour at room temperature covered from light on shaker. Plates were washed with 1X PBS and stained for DAPI (62248, Thermo Fisher Scientific) for one minute. Immunofluorescence images were visualized with an AMG EVOS FI AMF-4302 fluorescent microscope.

| Target | Source | Application |
| --- | --- | --- |
| Anti-53BP1 | Primary, Novus Bio, #NB100-304, rabbit | IF |
| Anti-phospho-Histone H2A.X (Ser139), clone JBW301 | Primary, Millipore, #05-636, mouse | IF |
| Anti-β-Tubulin III | Primary, Millipore, #T8578,<br>mouse | IF |
| Alexa Fluor 488 rabbit anti-mouse IgG (H+L) | Secondary, Invitrogen, #A-11059, rabbit | IF |
| Alexa Fluor 555 donkey anti-rabbit IgG (H+L) | Secondary, Invitrogen, #A-31572, donkey | IF |

### Neurite outgrowth

iNeurons were fixed and stained with β-Tubulin III for imaging. Images were obtained using an AMG EVOS FI AMF-4302 fluorescent microscope. Images were analyzed and quantified using ImageJ (National Institutes of Health). Neurite tracing was accomplished by inverting the image to a negative, setting a scale appropriate to the image objective, and manually tracing the length of the extension starting from the cell body and ending at the edge of the extension. Neurites per cell were quantified as a byproduct of neurite tracing. Neurite lengths were grouped by cell, denoting how many neurites originated from the same cell.

## Acknowledgments

This work was supported by grants from the National Institutes of Health, AG071815 to CS, The Audrey Myer Mars Foundation.

## Author Contributions

Performed experiments: SK, LA, NG, CS; Experimental design and interpretation of data: SK, AB, QL, EN, CS; Manuscript writing; SK, CS

## Competing Interest Statement

CS is Co-Founder of Hayflick Therapeutics. The authors declare no competing interests

---

**Figure S1.**
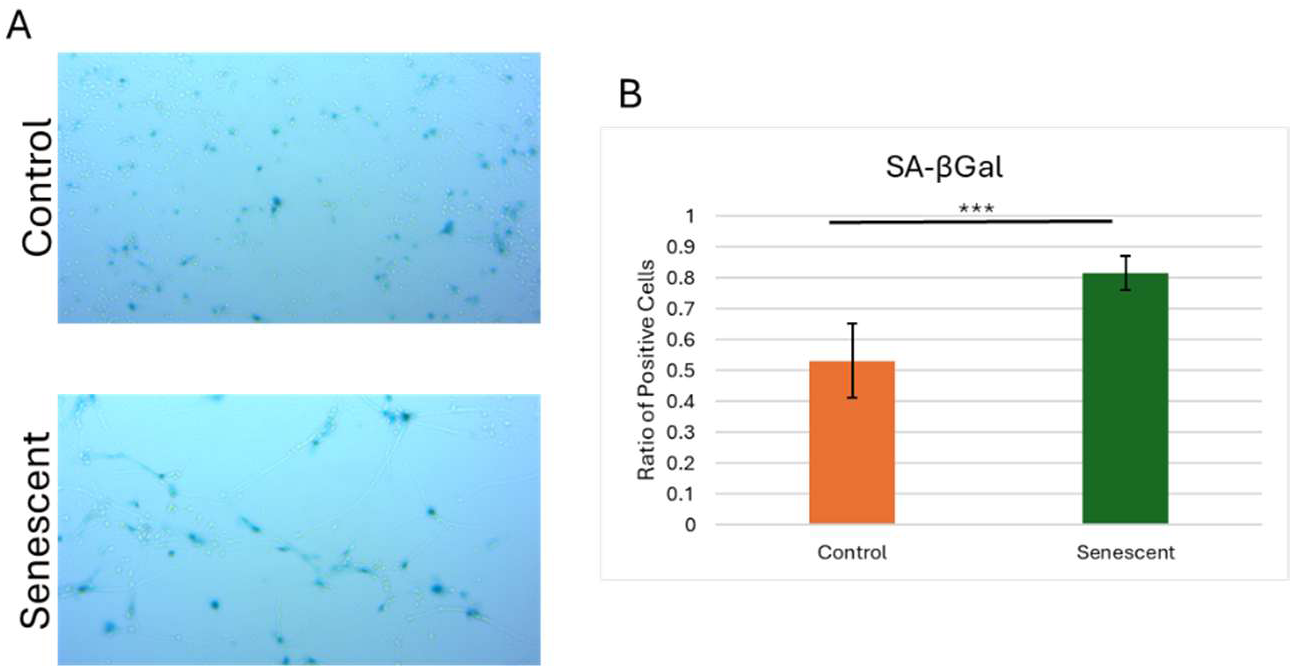
Etoposide treated astrocytes display increased staining for Senescence Associated Beta-galactosidase activity. Panel A contains representative image of control and senescent cells stained for SA-βGal and B contains quantification of the ratio of SA-βGal-positively stained cells.

**Figure S2:**
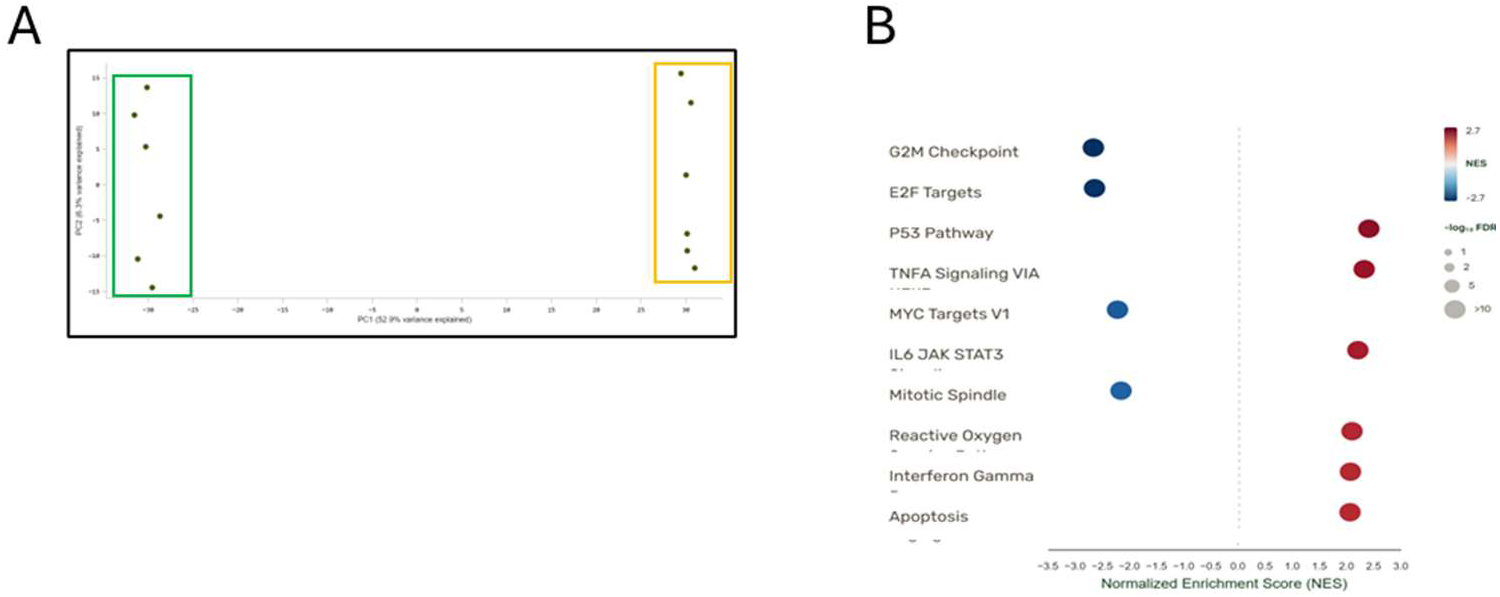
Transcriptional profile of senescent astrocytes. Panel A contains a PCA plot of RNA-seq profile. Green box highlights ETO-treated group and orange box highlights control group. Panel B contains gene ontology analysis for differentially expressed gene between senescent (ETO treated) and control astrocyte culture.

**Figure S3:**
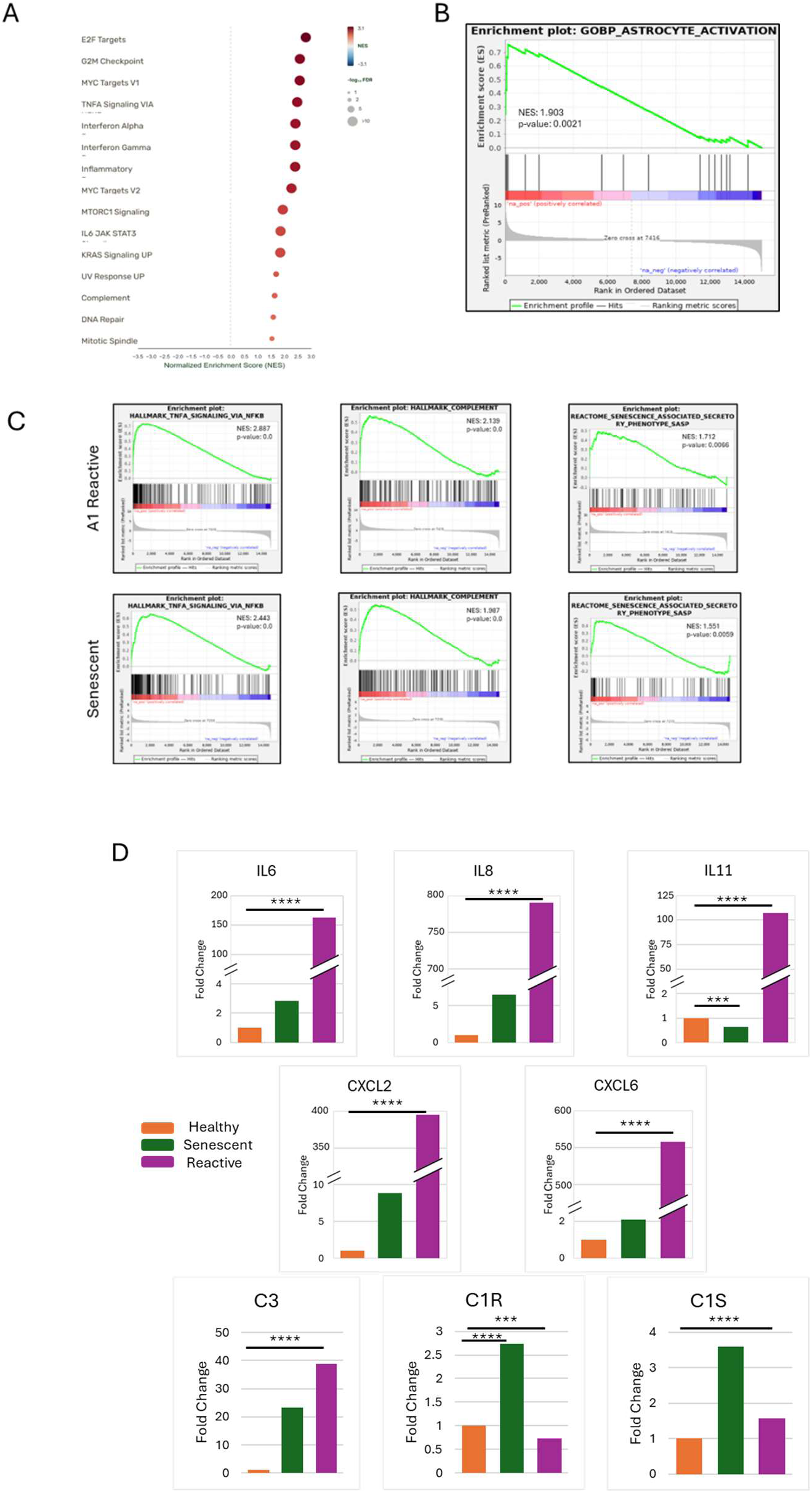
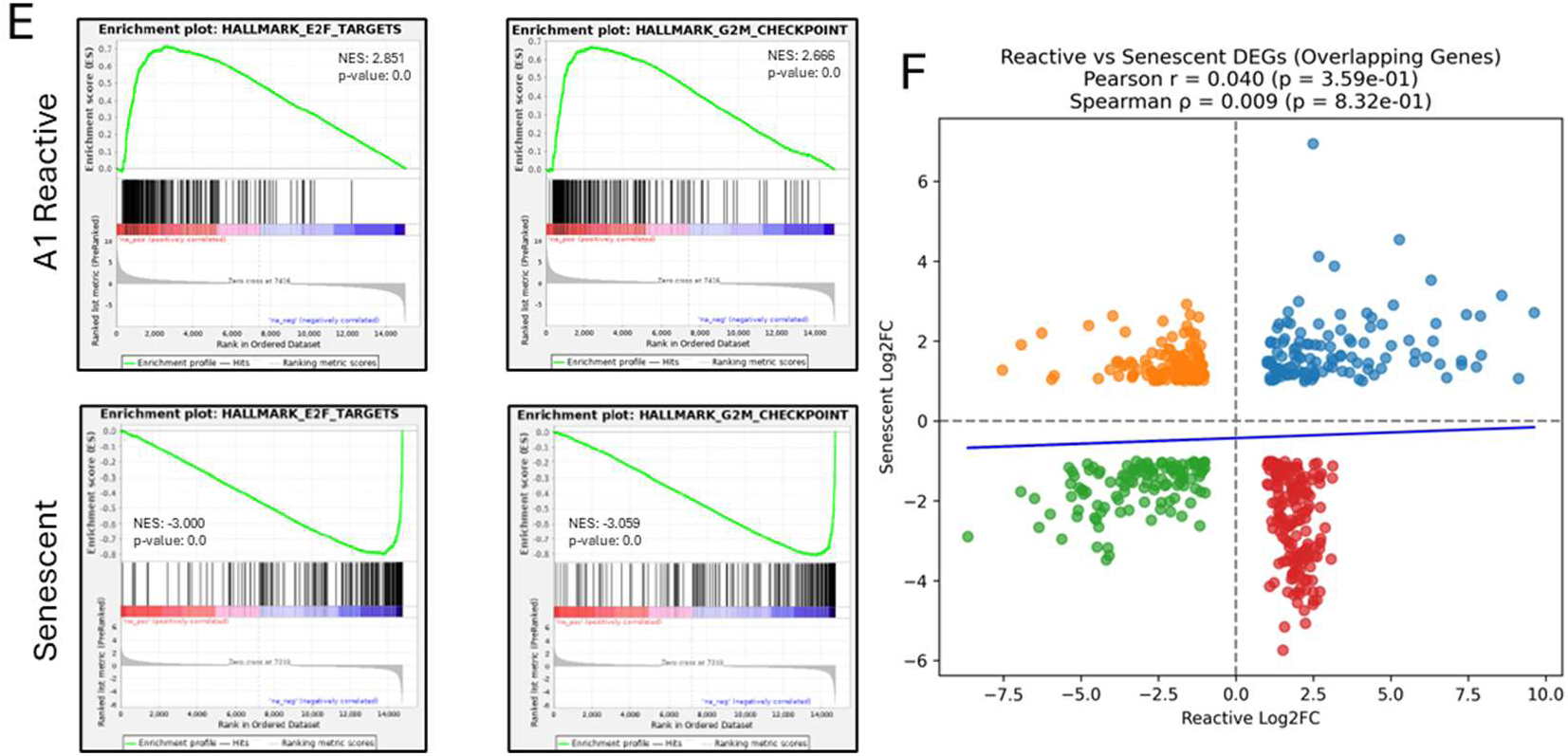
Reactive astrocytes display a distinct transcriptional profile from senescent astrocytes. Panel A contains a gene ontology analysis indicating increased expression of genes associated with proliferation (G2M, E2F targets) as well as inflammatory pathways (TNF, Interferon alpha and gamma, IL6 signaling) Panel B contains a GSEA analysis of reactive astrocytes confirming significant enrichment for reactive gene set while Panel C contains reactive and senescent transcriptomes against TNFα signaling via NFκB, Complement, and SASP pathways. Panel D contains direct count comparisons between senescent and reactive astrocytes for inflammatory cytokines (IL6, 8, 11, CXCL 2, 6) and complement factors (C3, C1R, C1S). Panel E contains a comparison of GSEA analysis for G2M and E2F targets for reactive and senescent astrocytes. Panel F contains a correlation analysis of human A1 reactive and senescent astrocytes. Statistical correlation was calculated utilizing both Pearson and Spearman analyses.

**Figure S4.**
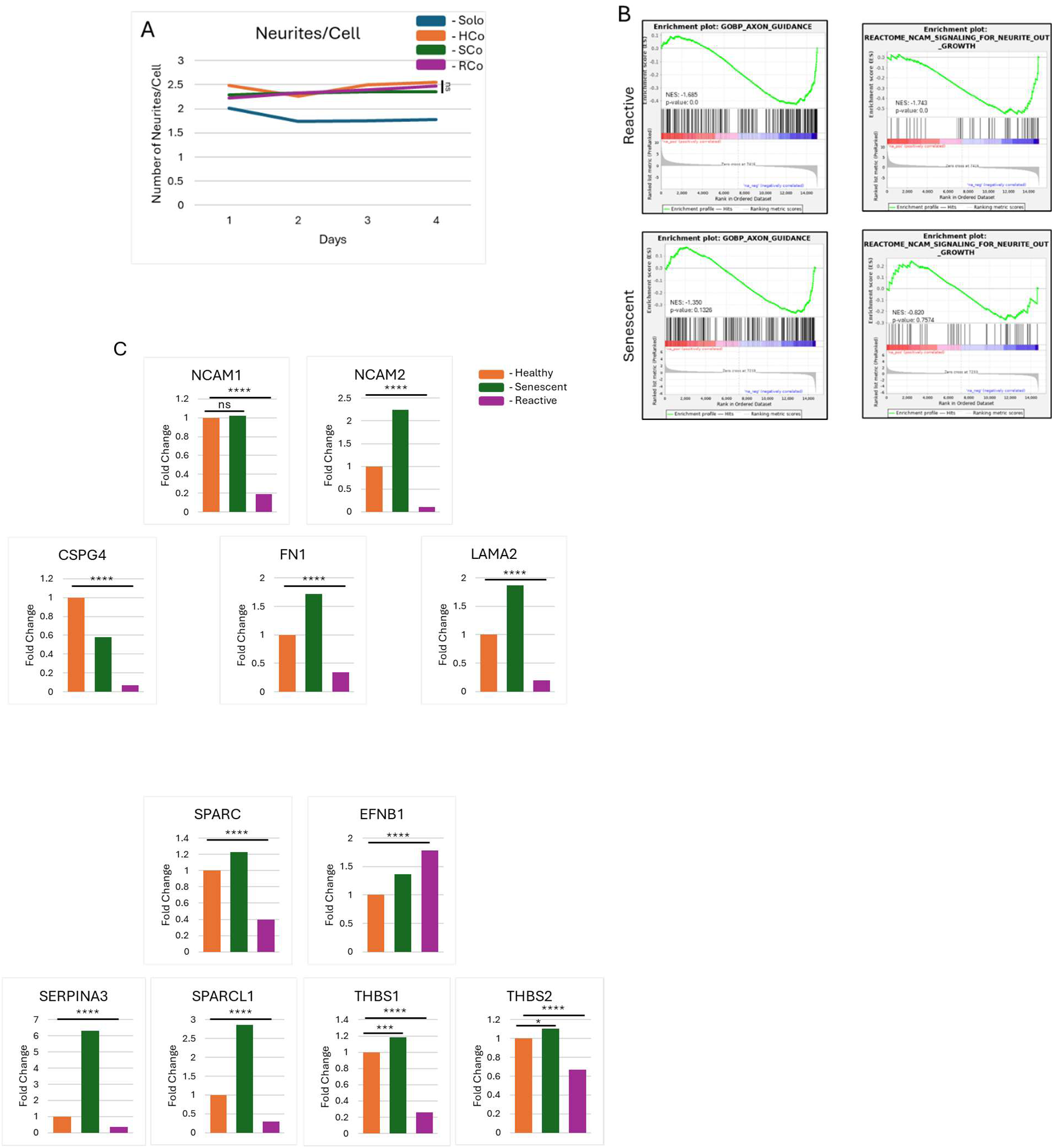
mRNA levels for genes associated with neurite outgrowth are reduced in reactive astrocytes. A) Average number of neurites per cell over 4 days. Panel B contains GSEA analysis for an axonal guidance and NCAM2 gene sets for reactive and senescent astrocytes. Panel C contains fold change comparisons for genes involved in neurite outgrowth, NCAM1, NCAM 2, CSPG4, FN1, LAMA2, SPARC, EFNB1, SERPINA3, SPARCL1, THBS1, and THBS2. Statistical analysis for neutite measurements were performed using a two-tailed unpaired student’s t-test and Mann-Whitney test with significance levels marked as *p < 0.05, **p < 0.01, ***p < 0.001, and ****p < 0.0001

**Figure S5:**
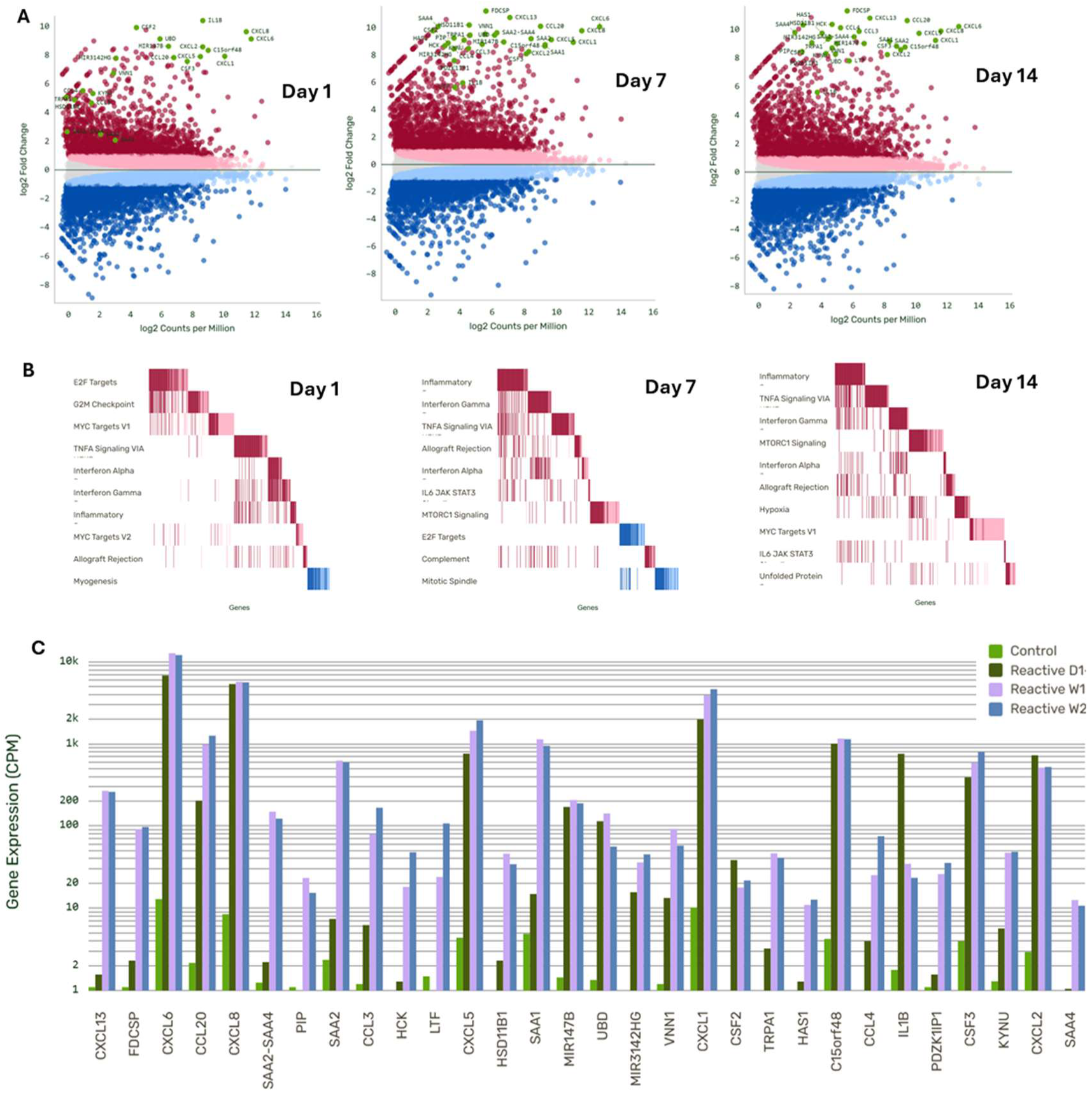
Reactive astrocytes maintain an inflammatory phenotype over time. Expression profiling for reactive astrocytes at days 1, 7, and 14 was carried out. Panel A contains volcano plots at day 1, 7, and 14. Panel B contains gene ontology analysis between control and reactive asctrocytes at day 1, 7, and 14. Panel C contains direct comparison of cpm values for the top 30 differentially expressed genes at day 1, 7, and 14.

## Notes

### Competing Interest Statement

CS is a Co-Founder and equity holder in Hayflick Therapeutics. This association has no influence on the work presented in this manuscript.

